# Rewiring of genetic networks in response to modification of genetic background

**DOI:** 10.1101/007229

**Authors:** Djordje Bajić, Clara Moreno, Juan F. Poyatos

## Abstract

Genome-scale genetic interaction networks are progressively contributing to map the molecular circuitry that determines cellular behaviour. To what extent this mapping changes in response to different environmental or genetic conditions is however largely unknown. Here we assembled a genetic network using an *in silico* model of metabolism in yeast to explicitly ask how separate genetic backgrounds alter network structure. Backgrounds defined by single deletions of metabolically active enzymes induce strong rewiring when the deletion corresponds to a catabolic gene, evidencing a broad redistribution of fluxes to alternative pathways. We also show how change is more pronounced in interactions linking genes in distinct functional modules, and in those connections that present weak epistasis. These patterns reflect overall the distributed robustness of catabolism. In a second class of genetic backgrounds, in which a number of neutral mutations accumulate, we dominantly observe modifications in the negative interactions that together with an increase in the number of essential genes indicate a global reduction in buffering. Notably, neutral trajectories that originate considerable changes in the wild-type network comprise mutations that diminished the environmental plasticity of the corresponding metabolism, what emphasizes a mechanistic integration of genetic and environmental buffering. More generally, our work demonstrates how the specific mechanistic causes of robustness influence the architecture of multiconditional genetic interaction maps.

## Introduction

Gene action is commonly determined by its interactions with other genes. This includes genes known to be associated to the action under study, but also those whose association is less expected or their biochemical properties still unknown. Both classes of interactions can now be effectively mapped at a large scale by following two complementary strategies.

The first one relies on the progress of experimental tools to produce genetic perturbations in large numbers and to automatically quantify their effects [1] (growth being typically the primary phenotypic readout but see, for instance, [2, 3]). These tools are now providing initial genetic landscapes of cells, e.g., [4, 5]. A second approach benefits from the advance of computational methods capable to predict phenotypes. Metabolic flux balance models are particularly useful in this regard, as they incorporate genomic information (of metabolism) into an *in silico* framework that can estimate cell growth under specific conditions [6]. Notably, flux balance predictions have been confirmed experimentally, e.g., [7]. Single mutant fitness and their corresponding genetic interactions can also be produced in this framework [8].

These strategies are currently being combined to better interpret the molecular underpinnings of genetic interactions (i.e., epistasis), both negative and positive. Negative epistasis (observed when the fitness defect of a double mutant is lower than that expected from single mutant values) indicates redundancy, that can reveal as functional associations between some pathways and/or complexes (e.g., the presence of negative epistasis between the urmylation pathway and the elongator complex in yeast suggested that both jointly modify certain transfer RNAs [5]), or as the buffering of alternative metabolic routes (e.g., leading to the synthesis of the same component [9]). Positive genetic interactions, in contrast, are commonly observed between genes that constitute a multi–protein complex or metabolic pathway, i.e., genes being part of the same functional unit [10]: a mutation in one of its constituents can inactivate this unit what reduces the effect of other perturbations in additional components.

Large scale approaches lead as well to the identification of system-level patterns, when the interactions are represented as genetic networks. For instance, the network presentation of high-throughput data of *Caenorhabditis elegans* and *Saccharomyces cerevisiae* clearly identified the presence of genetic hubs, that are mostly associated to chromatin regulation [5, 11]. Another feature, revealed for the first time with flux balance modeling, is monochromaticity; the specific distribution of epistasis types in the interactions within/between functional modules [12]. This characteristic was later confirmed by metabolic experiments [8] and high-throughput data [5], in which a specific distribution of epistasis strengths was additionally identified [13].

All previous properties implicitly suggest a stable architecture of genetic networks, a view that was partially influenced by the constant conditions in which interactions were examined. However, recent studies are emphasizing that this stability should not be necessarily the case. Genetic interactions and, more broadly, genetic networks were shown to change depending on the particular context where fitness is evaluated [10,14–17]. Rewiring is further confirmed by means of comparative analysis across organisms [4, 18–20]. Moreover, the “instability” of these networks should not come as a surprise; earlier works already discussed the influence of context (environmental, genetic) on the phenotypic effect of mutations and their interactions, e.g., [21, 22], a phenomenon that can directly influence evolutionary dynamics [23–25]. To what extent genetic networks are context-dependent is nevertheless mostly unknown.

Here we ask how the structure of a genetic network reorganizes in response to changes in gene background. To this aim, we mapped genetic interactions between metabolic genes by using a computational model of metabolism in *Saccharomyces cerevisiae*. The advantage of this approach (beyond avoiding experimental complexity) is that it enables the interpretation of the phenotype (i.e., fitness) as an univocal consequence of the structure of the metabolic reaction network underneath. We consider two broad (genetic) background classes. The first class corresponds to single gene deletions of each of the enzymes that are active (i.e., showed nonzero flux) in wild-type (WT) conditions. We characterized which type of backgrounds originate stronger network change, and in which kind of interactions this variation is more pronounced. The rewiring patterns found stress the different organization of biosynthetic and catabolic routes and how this impacts their capacity to compensate change. A second class presents neutral backgrounds, that are generated by a trajectory of accumulated neutral mutations. This helps us to appreciate how cryptic variability modifies buffering in genetic networks, and how the new network structures associate to differential environmental plasticity. We additionally corroborate some of these patterns with inspection of experimental data.

## Results

### Rearrangement of the catabolic repertoire determines genetic rewiring

We began analysing the rewiring observed in genetic backgrounds defined by deletions of single genes that are metabolically active in WT conditions (Materials and Methods). These deletions normally result in the blockade of some (nonessential) reactions and the corresponding reconfiguration of metabolic fluxes [9, 26]. How is this readjustment contributing to genetic rewiring?

To understand this correspondence, we quantified the number of modified interactions with respect to the WT genetic network as a simple rewiring score (Fig. 1.A), and also counted the quantity of altered fluxes as measure of the underlying metabolic readjustment. Fig. 1.B shows the strong relation between these two scores (Spearman’s *ρ* = 0.72, *p* < 10^−8^). We subsequently partitioned the metabolic measure into a qualitative and quantitative component (i.e., active fluxes that become inactive or vice versa, and changes in relative flux through already active reactions, respectively). Qualitative changes predict the strength of genetic rewiring (multiple linear regression *p* < 10^−8^) while quantitative ones do not (*p* = 0.97). Thus, genetic rewiring denotes a redistribution of metabolic fluxes to alternative reactions and pathways.

**Figure 1.**
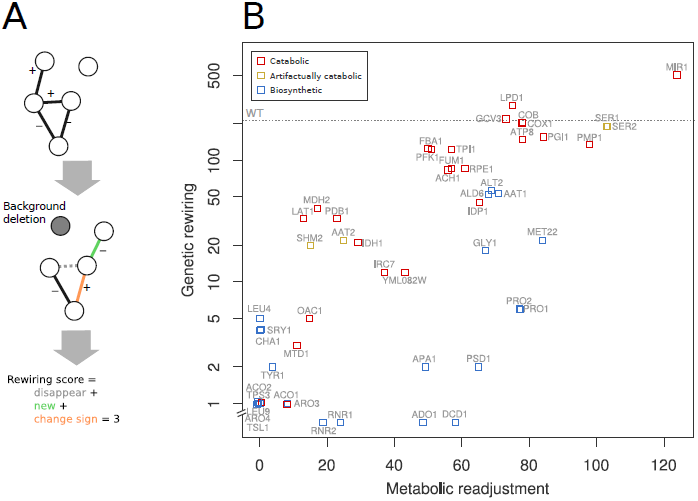
**Genetic rewiring correlates with the underlying metabolic readjustment.** (A) Cartoon that illustrates the kind of alterations in the genetic network that contribute to the rewiring score: new interactions + interactions that changed sign + disappeared links (excluding those of the gene acting as background). (B) Association between genetic rewiring and metabolic readjustment (as number of reactions that modified their relative flux) in response to deletions in active genes, i.e., backgrounds; there exists 207 interactions in the WT network (dashed line), and 277 active fluxes in the corresponding metabolism (coordinates incorporate some noise to help visualization, and the *y*-axis logarithmic scale is broken to locate backgrounds with no genetic rewiring). Notably, catabolic genes exhibit much stronger rewiring than biosynthetic ones, regardless of the strength of metabolic readjustment (some genes are artifactually catabolic in the model, see Supplement).

We then explored if the perturbation of any specific metabolic function leads to particularly strong rewiring. If backgrounds linked to a given function result in strong rewiring, this should be detectable as variation on its associated metabolites. We found that rewiring is predicted by the variation of several metabolites whose production and homeostasis is tightly connected to catabolism (e.g., phosphate, redox equivalents, quinones, or intermediaries from glycolysis or TCA cycle, Supplement and Table S1). Indeed, backgrounds corresponding to catabolic functions display stronger rewiring than those related to biosynthetic functions (Fig. 1.B). Note also that the strong genetic rewiring produced by some apparently biosynthetic genes (e.g., SER1, SER2 or SHM2) revealed upon detailed inspection an artifactually catabolic role of these genes (e.g., ATP synthesis, Supplement and Table S2) that validates a general catabolic underpinning of network rewiring.

### Rewiring is pronounced between functional modules and in weak genetic interactions

We next examined the association of rewiring with the underlying metabolic organization and the type of genetic interaction. We initially computed the instability of each WT interaction by simply counting the number of backgrounds in which it changes (from a total of 100). Interactions tend to be conserved with instability being stronger for those constituted by genes belonging to different metabolic annotation groups (mean instability within same group = 5.1, mean between groups = 7.8, *p* = 3.4 × 10^−5^). New interactions exhibit as well a tendency to connect different modules (90% of cases) compared to the WT links (78%, Fisher’s test *p* < 10^−7^). This pattern resembles recent reports that detected stronger instability in interactions established between (rather than within) functional modules, but in response to environmental change [15, 16].

Fig. 2A explicitly illustrates the concentration of genetic interactions in catabolic modules (Table S3) as well as their high relative instability. Conversely, biosynthetic modules exhibit fewer and more stable links. Catabolic modules are characterized in addition by the appearance of new interactions in different backgrounds. In contrast, new links arise less frequently between biosynthetic modules, and in a more background-specific manner (Fig. 2B).

**Figure 2.**
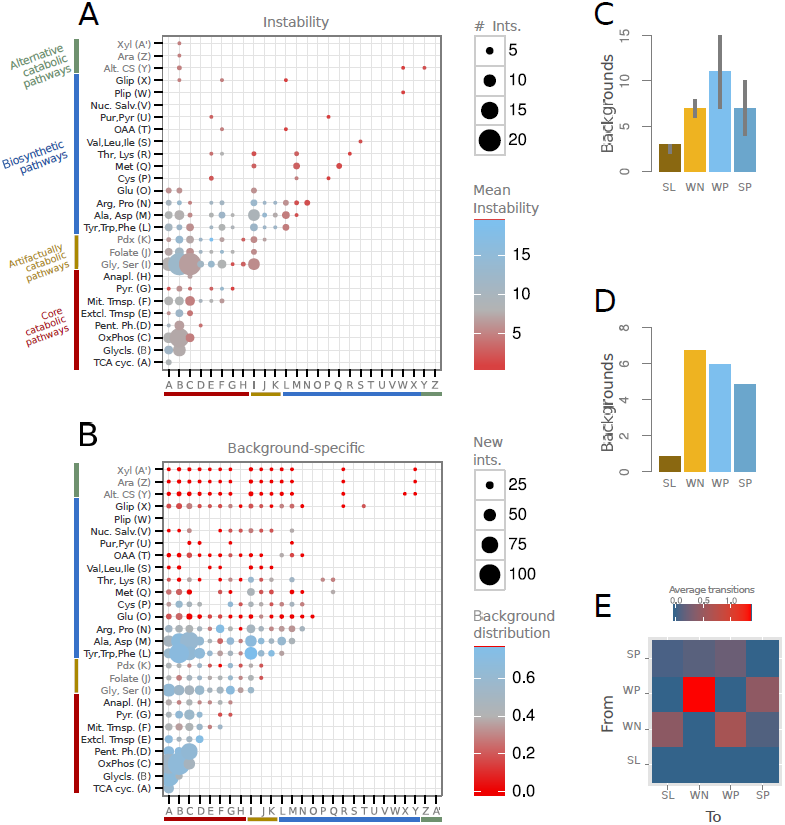
**Catabolic and biosynthetic modules exhibit distinctive genetic rewiring.** (A) Number of interactions of the WT network between the corresponding metabolic annotation modules (dot size), and associated average instability (dot color; measured for each interaction as the number of backgrounds where it disappears, changes sign or strength). Catabolic modules show higher instability than biosynthetic ones. (B) New interactions (i.e., absent in WT network) between modules (dot size), and their distribution among backgrounds (dot color; distribution quantified as normalized Shannon entropy, Materials and Methods). Catabolic modules are characterized by the emergence of many new interactions in different backgrounds. Fewer new interactions appear among biosynthetic modules, these being generally more background specific. (C) Instability of interactions as a function of sign and strength (median with upper and lower quartiles). Weak interactions are more unstable than strong ones of the same sign, and positive interactions are more unstable than negative. Similarly, in (D) we show how WN interactions appear dominantly in a new background, on average. (E) Expected number of transitions between interaction types in an average background. WP to WN conversions are the most recurrent ones. Interaction classes are synthetic lethal (SL), weak negative (WN), weak positive (WP), and strong positive (SP), see Material and Methods.

Moreover, the strength and sign of the interaction influences as well its instability. We observed weak interactions to be more unstable for each epistasis type (Fig. 2C). While this could be associated to the fact that weak links tend to appear between module (Fisher’s test, *p* < 0.0002), both being weak and between-module independently correlate to instability (Supplement, Fig. S1, Table S4). Within new interactions, we also noticed that weak negative (WN) are the dominant class (Fig. 2D) and the ones that take part of most sign changes (which usually occur between WN and weak positive, WP, Fig. 2E).

### The structure of rewiring evidences the intertwined organization of catabolism

Deletion of catabolic genes originates therefore a strong rewiring of genetic interactions (Fig. 1). That these genes exhibit large connectivity (genetic hubs correspond almost uniquely to catabolic genes, Fig S2), and that this connectivity is dominantly constituted by weak interactions (i.e., nodes with weak average epistasis |*ε*| < 0.5 present a mean of ∼8 connections, while those with strong average epistasis, |*ε*| > 0.5, present ∼3, Wilcoxon-test *p* = 0.003) suggests that the response to a change in (catabolic) backgrounds is very much linked to the rewiring of hubs –recall the stronger instability of weak links between catabolic modules, Fig. 2– what ultimately indicates the intertwined organization of catabolism (Fig. 3).

**Figure 3.**
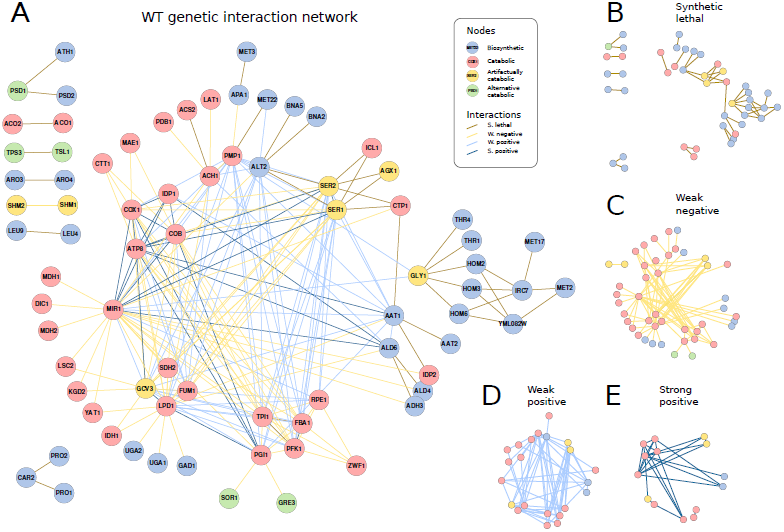
**The topology of the WT genetic network reflects the intertwined organization of catabolism.** (A) WT genetic network with nodes colored by function and interactions by class. (B)–(E) Decomposition of the network into the four interaction classes. Note the association between these types and metabolic function (i.e., synthetic lethal links broadly corresponding to biosynthetic genes and the rest –weak negative, weak positive, strong positive– to catabolic) that we additionally corroborated with experimental data of epistasis between metabolic gene pairs [8] (Fig. S24). The distribution of weak interactions emphasizes the intertwined organization of the catabolic core (some genes appear as artifactually catabolic in the model, see discussion in Supplement). Moreover, synthetic lethal links appear in the “periphery” of this core (full details of each synthetic lethal cluster in Supplement, Figs. S27–S33).

This organization is effectively described by the action of a number of versatile genes capable to contribute to fitness in different ways (by altering their function) and to partially buffer each other’s action (Fig. 3C-E). The abundance of WP interactions emphasizes the different means to contribute to fitness. For example, although glycolysis (e.g., genes such as TPI1, FBA1) and the TCA cycle (LPD1, FUM1) work in coordination to supply reduced equivalents to oxidative phosphorylation, they can also readjust their metabolic role when one of the subsystems is compromised (what causes WP links between TPI1 and LPD1 or FBA1 and FUM1; Fig. 3D). Moreover, WN interactions indicate distributed buffering [27] among sets of genes that can implement a particular metabolic action with different degree of efficiency (Fig. 3C). An example corresponds to MIR1 that negatively interacts with a group of genes that jointly represent a metabolic alternative for mitochondrial phosphate import (Supplement, Figs. S3–S5). These include genes that contribute to the phosphate import strictly speaking, but also many that reduce the redox imbalances generated by this alternative mechanism (as a result of malate being antiported out of mitochondria in exchange for phosphate), with far reaching consequences regarding the whole set of fluxes through catabolic pathways.

In Fig. 4A, we showed in detail an example of the rewiring experienced by a hub (PFK1) in different backgrounds (Figs. S6–S14 for rest of hubs). Despite the instability of their interactions, hubs tend to exhibit a remarkable connectivity conservation (Fig. S15). We further display the rewiring caused by PFK1 acting as background (Fig. 4B), and how hubs principally rewire their interactions in response to the deletion of other hubs (Fig. 4.C); this demonstrates the functionally intertwined architecture of catabolism. Notably, we detected a number of genes that acquire the role of genetic hubs in some specific contexts, and that frequently corresponds again to catabolic functions (Fig. 4.D).

**Figure 4.**
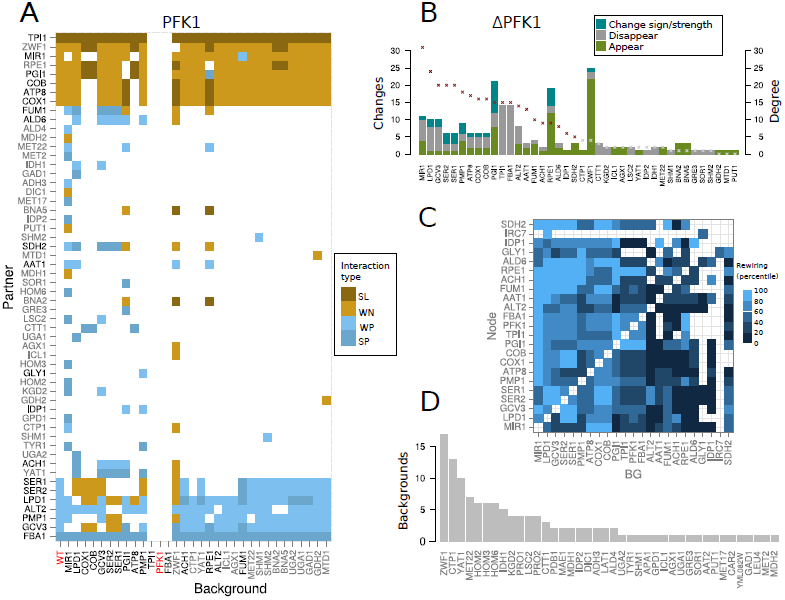
**Rewiring of hubs.** (A) Map of the genetic rewiring of interactions between hub PFK1 and its partners (vertical) in all backgrounds where rewiring of this hub is observed (horizontal). First column corresponds to the WT. Names of backgrounds and partners that are also genetic hubs are highlighted in black. PFK1 shows no connectivity in gene contexts ΔFBA1, that presents a SP link with PFK1, and ΔTPI1, that presents a synthetic lethal link (i.e., PFK1 becomes essential in these backgrounds). Interaction classes denoted as Fig. 2. (B) Rewiring experienced by genes in a ΔPFK1 background (color coding illustrates the different types of rewiring). Note that most of the rewiring in this background is observed in hubs (gene connectivity, i.e., degree, also shown) –most specially, in those that are functionally related, e.g., PGI1, TPI1, FBA1– and pentose phosphate genes that assume initial glucose processing after PFK1 deletion. (C) Rewiring of hubs in response to hub deletions. Each rewiring score was normalized by the connectivity of the specific hub in the WT network. The map represents the percentiles of these normalized scores [with percentiles (20, 40, 60, 80, 100) corresponding to values of (0.22, 0.40, 0.66, 0.96 and 5.6); a value of 1 means that the number of rewired interactions equals WT connectivity; empty spaces correspond to absence of rewiring]. The only hub with a purely biosynthetic function (IRC7) is the one deviating most for this general pattern. (D) List of genes acting as condition-dependent hubs, and number of backgrounds in which they exhibit such large connectivity; many of these genes are related to catabolism.

### Genetic rewiring in neutral backgrounds indicates reduction in buffering

We now introduce a second class of genetic backgrounds in which a set of neutral gene deletions accumulate (Materials and Methods, and Fig. 5A for a case trajectory). How does the network rewire in response to these backgrounds? We answer this question by discussing first which genes appear in neutral backgrounds and how their mutation modify the network.

**Figure 5.**
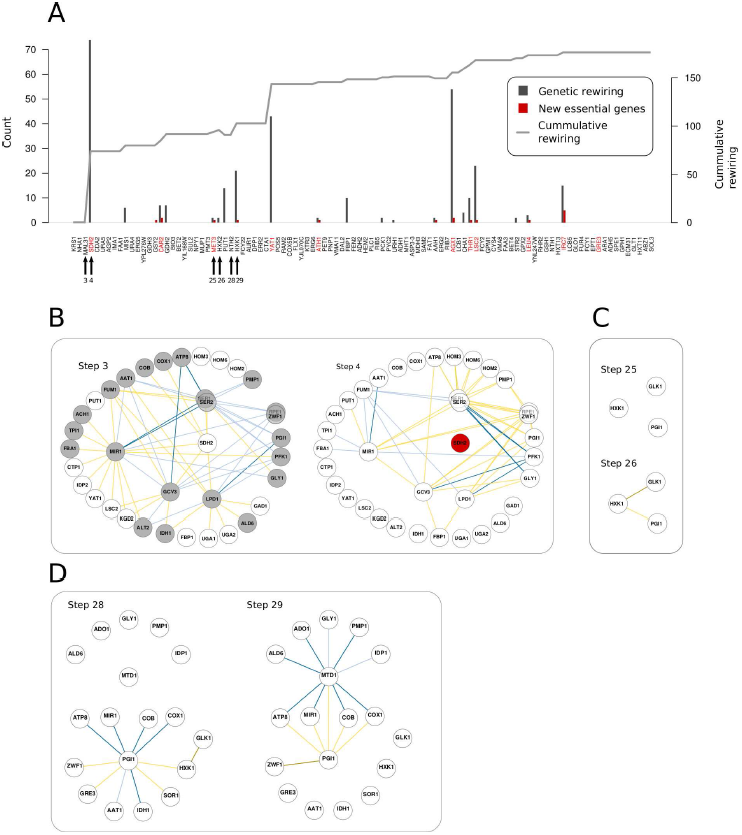
**Detailed view of a neutral deletion trajectory.** (A) Genes were consecutively removed (from left to right; nodes of the WT network in red) what accumulates network rewiring (curve). Bars depict “local” change or rewiring score and number of essential genes (both with respect to the previous step). Many deletions do not affect network topology but some few ones produce strong rewiring. The appearance of essential genes is concomitant with variable amounts of genetic rewiring. (B) to (D) show the sub-networks that get to be modified before and after several critical steps of the trajectory. Deletion of SDH2 (B) reveals second-order alternatives to the action of this gene in the “buffering ladder” (as new negative interactions). Several genes additionally undergo changes in their functional role that are reflected in modification of their interaction signs and, notably, in the emergence of new positive interactions. In (C), after the deletion of one of the three alternatives for hexokinase (HXK2), the other two constitute a synthetic lethal link (HXK1 and GLK1). HXK1 is also able to phosphorylate fructose and this underlies a supplementary weak negative interaction with PGI1. Finally, in (D), deletion of HXK1 further rewires PGI1 interactions (e.g., the weak negative link with pentose phosphate pathway gene ZWF1 becomes synthetic lethal, as the only alternative for initial glucose processing) and induces several positive links at a relatively distant part of catabolism, e.g., MTD1, involved in the artifactual glycine fermentative pathway (color code of genetic interactions as previous figures).

Note that a subset of these deletions could correspond to nodes of the WT network. This implies genes that are part of negative interactions, and broadly causes the emergence of novel buffering connections (in terms of new interactions or change of sign of WT links). This is observed for instance in the deletion of SDH2 in Fig. 5A, a gene that acts as buffer in WT conditions of GCV3, LPD1, etc (Fig. 5B). SDH2 deletion uncovers secondary buffering mechanisms (e.g., by FBP1 or GLY1). Changes of sign in WT links are observed as well; for example, interactions with MIR1 or ZWF1. These changes are typically exhibited by catabolic genes in which different pathways often contribute to fitness either feeding one another linearly (e.g., to ultimately supply redox power to oxidative phosphorylation) or serving as substitutes to each other (e.g., as direct mechanisms of ATP synthesis). In some cases, the new buffering mechanisms can additionally modify how active genes contribute to fitness by removing, or adding, positive interactions to the network (e.g., ALT2–GCV3 link in Fig. 5B). Note also how genes that become nodes in one step of the trajectory can produce relatively strong rewiring when they undergo subsequent deletion (e.g., FBP1, that turns into a node after SDH2 mutation to be deleted in a later step, Fig. 5A).

Other genes that arise in neutral backgrounds (not being network nodes) can reduce too buffering alternatives. This can be exemplified, for instance, with the deletions of MIS1 or HXK1/HXK2 in Fig. 5A. The latter is especially illustrative, as three different genes are capable of performing the glucose phosphorylation reaction: HXK1, HXK2 and GLK1. HXK2 is deleted first, and the two remaining ones then exhibit a SL link (Fig. 5C). When HXK1 is later deleted, GLK1 becomes essential. However, HXK2 and HXK1, but not GLK1, are capable to perform additional functions as hexokinases, such as fructose or mannose activation. Although not essential, reduction of these functions induces many new catabolic constraints that uncover new genetic interactions: negative ones, between glycolysis and respiration, and positive ones, between respiration and folate pathway genes (Fig. 5D).

Similar analyses of a group of 200 alternative neutral backgrounds contribute to distinguish common patterns of change. First, the corresponding networks exhibit a smaller number of nodes (196 out of 200 contain less nodes than WT). This signal relates of course to the deletion of negatively interacting nodes (Fig. S16) that points to modifications of buffering mechanisms. Second, networks tend to exhibit more epistatic interactions per node (166 out of 200 present larger average epistasis), a pattern again related to negative interactions (Fig. S16). Third, the negative component of the network undergoes a significantly stronger rewiring than the positive one (Fig. S17). This suggests overall that previously hidden phenotypic effects unveiled as a result of the global reduction in buffering mechanisms. This is further evidenced by an increase in the number of essential genes, which is observed in 93.5% (187/200) of genotypes and occurs concomitantly with rewiring of the network.

### Catabolic perturbations associate network instability to diminished environmental plasticity

One could expect that not all accumulated deletions in the neutral backgrounds impair metabolism in the same way, and thus different backgrounds could cause contrasting metabolic plasticity. This can be detected by generating diverse random environments [28] (Materials and Methods) in which fitness is computed. The resulting growth measures did reveal the cryptic variability linked to the neutral trajectories (Fig. 6A).

**Figure 6.**
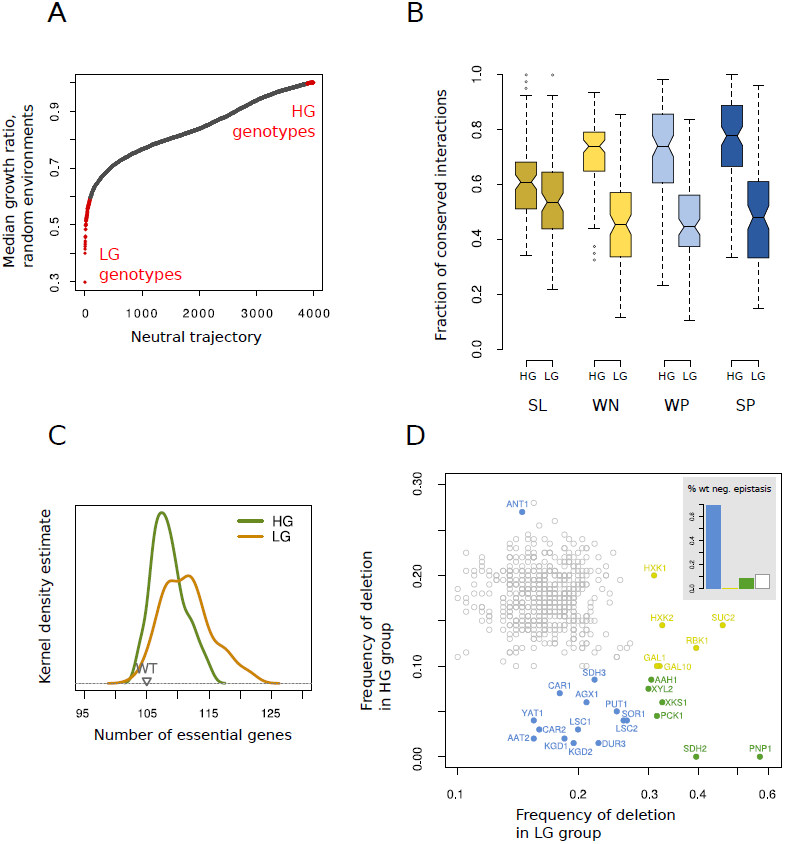
**Genetic rewiring patterns predict loss of environmental plasticity in response to neutral backgrounds.** (A) Environmental plasticity exhibited by 4000 different metabolisms originated from the WT after following a neutral deletion trajectory. Plasticity is scored as the median of the fitnesses (growth ratio) of each metabolism in 1000 randomly generated environments (growth ratio, in a given environment, is the division between growth rate of a particular metabolism to the WT growth rate). We highlight in red the groups with *highest* and *lowest* growth (HG and LG, respectively). (B) Fraction of conserved interactions by type and LG or HG genotypes. (C) Distribution of the number of essential genes in the HG and LG groups. (D) We performed a bootstrapping analysis to check for enrichment/depletion of each metabolic gene in the LG or HG trajectories. Genes significantly enriched (or depleted) in the HG genotype, LG genotype or both were correspondingly colored in blue, yellow and green (*p* < 0.01, after multiple testing). Inset. Percentage of genes with genetic interactions in the WT network in these significant groups. Interaction classes denoted as Fig. 2.

To relate this variability to specific rewiring patterns (in WT conditions), we assembled the genetic networks of the 100 genotypes with the highest (and lowest) median growth (HG and LG, respectively; note that these are the 200 networks considered as a whole before). Those networks corresponding to LG genotypes display a significantly stronger rewiring (Fig. 6B and Fig. S18), and are considerably smaller both in number of nodes and edges (Fig. S19). In addition, interactions associated to catabolic pathways were considerably less conserved in LG genotypes (Fig. S20).

Although the number of new interactions did not differ between LG and HG (mean = 60.6 and 61.0 respectively, *p* = 0.96, see also Fig. S21A), a subset of them appeared more frequently in LG genotypes–notably, negative ones between pentose phosphate pathway (ZWF1 and RPE1) and other catabolic genes (e.g., PGI1, MIR1 or LPD1, Fig. S21B-C). Sign and/or strength change was also considerably stronger in LG genotypes (mean = 24.3 interactions/genotype) as compared to HG (mean = 13.4 interactions/genotype, *p* = 6.7 × 10^−14^, Fig. S21D-E). As a result of the stronger rewiring, LG networks exhibited higher negative-to-positive interaction ratios than HG ones (mean = 1.65 *vs.* 1.53, Wilcoxon’s test *p* = 0.0005). The number of essential genes was higher as well (Fig. 6C), including several crucial catabolic components (e.g., ATP8, FBA1, PGI1, Fig. S22).

These results evidence that the loss of catabolic buffering mechanisms underlies both genetic network rewiring and reduction of environmental plasticity in LG genotypes. Namely, carbon sources other than glucose usually require only few transformation steps before being incorporated into the core catabolic pathways, e.g., at different steps of glycolysis or TCA cycle. Some other sources are alternatively transformed into glucose by means of gluconeogenesis. Core catabolic pathways are used then with relative independence of the external carbon source. However, they can be used differently: some branches that are optimal in one environment can be suboptimal in another (where they can nevertheless serve as an alternative to the optimal one). This is further corroborated by the differential distribution of deletions between LG and HG genotypes (Fig. 6D). LG genotypes are enriched in 26 specific deletions that can be grouped in i/ genes important for the initial processing of different carbon sources (e.g., PNP1, XYL2, XKS1, GAL1, etc), ii/ gluconeogenesis (e.g., PCK1), and iii/ key catabolic genes, such as SDH2, KGD1, or LSC1. Although neutral in glucose minimal medium, they constitute buffering mechanisms for deletions in other catabolic genes (evidenced also by their multiple negative interactions) but importantly can take over their role under different carbon sources.

## Discussion

Different genetic backgrounds can modify gene interactions and consequently rewire genetic networks. Here, we systematically examined how backgrounds impact networks by using an *in silico* model of yeast metabolism [6] (Materials and Methods).

We first analyzed to what extent strong genetic rewiring is necessarily coupled to strong metabolic readjustments. To this aim, we introduced a class of backgrounds defined by single deletions of active enzymes (i.e., that exhibit nonzero flux in WT conditions). As simple score to measure metabolic reorganization, we counted the number of reactions with a change of flux upon deletion. We found a relatively large number of backgrounds that exhibit substantial readjustment. However, only those that involved a switch to alternative metabolic pathways appeared coupled to strong genetic rewiring (Fig. 1). These backgrounds correspond to a set of catabolic genes that act as genetic hubs (Fig. 1 and Fig. S2). Interestingly, we also noticed strong genetic rewiring in genes that (artifactually) act as catabolic in the *in silico* yeast (see Supplement) which further confirms the linkage of metabolic readjustment and strong rewiring through catabolism.

In fact, most of the genetic instability is observed in the densely connected sub-network associated to central catabolic functions, which is mainly composed of positive and weak interactions between different functional modules (Figs. 2–3). In this way the structure of the wild-type network is already representing the functional associations that are more sensitive to background change. In addition, most of the background-specific (i.e., not observed in the WT) interactions and hubs are as well related to catabolism and enriched by intermodule weak negative epistasis (Fig. 2B and Fig. 4D).

The strong correlation between rewiring and perturbation of currency metabolite balances (Supplement, Table S1) greatly helps to understand these patterns. The existence of multiple NADH/NADPH and ATP producing enzymes in catabolic pathways enables them as potential substitutes to each other, i.e., as regulators of currency metabolite homeostasis [29]. But these mechanisms are not equivalent biochemically and consequently not equally optimal (Supplement and Figs. S3–S5 for an example). Catabolism exhibits then a functional degeneracy in which qualitatively different catabolic configurations lead to similar but not identically efficient solutions [27]. This explains many of the phenotypic features corresponding to catabolic genes. In particular, it causes catabolic genes to be typically nonessential, but often fitness contributing. Degeneracy explains too the pervasiveness of weak and unstable interactions which reflects either fitness contributions that are partially shared (weak positive), or deficient buffering (weak negative). Notably, background-dependent interactions tend to be weak as well. In sum, the distributed nature of catabolic processing determines the transient and context-dependent functional associations that define its epistasis network.

Biosynthetic pathways exhibit in contrast a much different architecture. They usually display relatively isolated and linear configurations, each of them containing very specific metabolites that simply act as intermediates for the synthesis of a particular compound. This limits the buffering possibilities compared to catabolism what is manifested in the enrichment of essential genes, and also in a genetic landscape dominated by redundancy-based synthetic lethal interactions (Fig. 3, Supplement and Fig. S27–S33). These SL interactions form smaller (peripheral) clusters and exhibit a marked stability that only becomes disrupted when one of the partners is deleted or becomes essential (Figs. 2–4).

These findings are consistent with several evidences from previous experimental studies on the rewiring of genetic interactions across species (studies not always linked to metabolism). For instance, SL pairs and interactions within functional modules were found considerably conserved, while interactions between modules remodeled [4, 18] – both signals confirming what we observed–, and the change of epistasis sign that we detected in catabolic nodes could indicate a sort of functional repurposing [19]. Moreover, we also recognized an association between interaction type and metabolic function (as we found *in silico*) in a set of genetic interactions recently measured experimentally between metabolic genes (Fig. S24) [8].

Likewise, our analysis clarifies the linkage among fitness contribution, pleiotropy, and network connectivity (node degree) [8]. That a particular gene is nonessential but contributes to fitness implies the existence of a number of inefficient distributed buffering mechanisms [9] of the type observed in catabolism. Pleiotropy is in addition strongly related to catabolism, due to the participation of currency metabolites in all biosynthetic routes of biomass constituents. Pleiotropy is thus present in catabolic genes and absent in biosynthetic ones (Table S5), and its correlation with fitness contribution and node degree could in the end denote the distributed robustness of the catabolic subsystem. As expected, both pleiotropy and fitness contribution anticipate rewiring (Fig. S23).

We examined a second class of backgrounds that are rather defined by (the accumulation of) neutral deletions (Fig. 5) [30, 31]. These trajectories generally originated metabolisms with a higher incidence of essential genes and smaller but more densely connected genetic networks (Fig. S16). This denotes overall a global reduction in buffering (Fig. S19). Neutral backgrounds also modify environmental plasticity (i.e., capacity for robust growth in a range of environments) in divergent ways (Fig. 6A–C and Figs. S20–S21). Notably, genetic networks associated to more limited plasticity present strongest genetic rewiring that it is mainly observed in interactions between genes associated to catabolic function (Fig. 6D). The mechanistic explanation is that, after usually few initial specific processing steps, all contrasting carbon sources enter the common catabolic core (glycolysis, TCA cycle, respiration). Mutations that are neutral in glucose minimal medium (affecting less efficient catabolic routes) can nevertheless represent the most efficient catabolic processing alternatives in other carbon sources.

The connection between environmental and genetic robustness [32] would further predict that the patterns identified in response to the alteration of genetic background could be similarly recognized in reaction to environmental change. To test this hypothesis, we characterized rewiring of a recently assembled (yeast) genetic network after several DNA-damaging treatments [17]. Conservation of the untreated network is well predicted by our model, with weak interactions being more unstable than strong, and positive more unstable than negative (Fig. 7A, compare with Fig. 2C). Interactions among genes that are functionally related were also more stable (Fig. 7B). Moreover, treatment-specific links occur between functionally different genes (92% as compared to 87% in the untreated network, *χ*^2^-test, *p* = 3 × 10^−15^) and are more often weak (Fig. S25).

**Figure 7.**
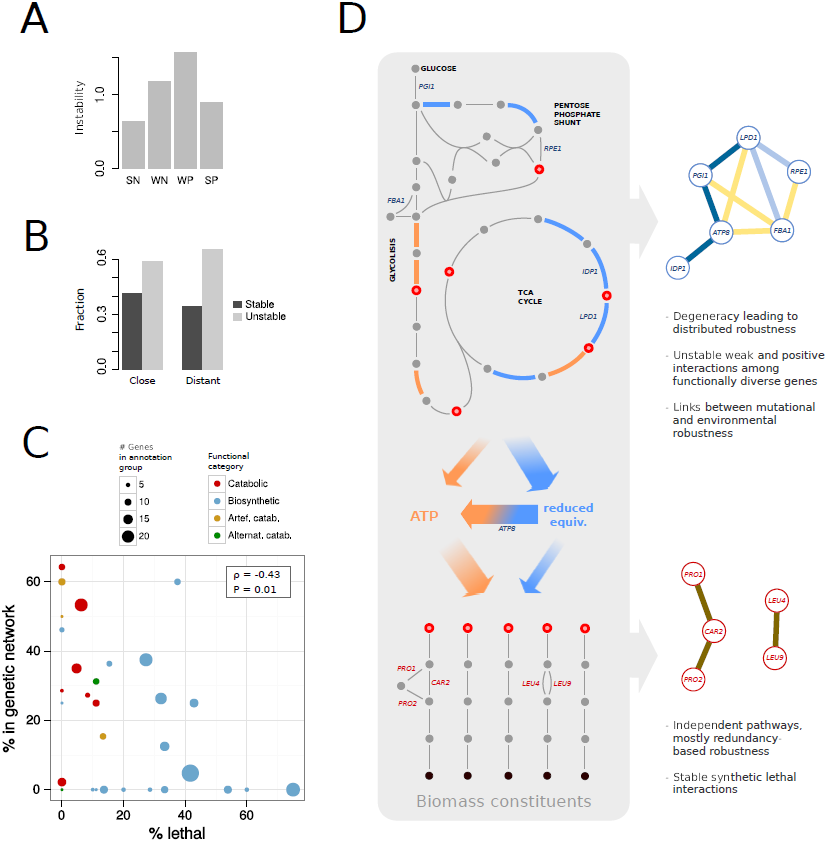
**Robustness influences rewiring patterns.** (A) Relative instability of each interaction type after environmental change (interactions were experimentally characterized in yeast growing in rich media and in three different environments [17], Material and Methods). This pattern is qualitatively similar to the one predicted in response to modifications of genetic background (Fig 2C). (B) We observe a higher proportion of unstable interactions among those genes functionally distant (*χ*^2^-test, *p* = 0.003, Materials and Methods). (C) Robustness and genetic landscape clearly distinguish catabolic from biosynthetic modules. Each dot represents a metabolic module with size and color indicating number of genes in the module and functional category, respectively. We show percentage of essential genes in each module (as a proxy of robustness; horizontal axis), and number of (nonessential) genes that are nodes in the WT genetic network (as a proxy of genetic landscape; vertical axis). We further confirmed this signal with experimental data (Materials and Methods) (D) The architecture of catabolism and biosynthesis (left) determines the resulting genetic network and its stability (right). We show in blue the reactions producing NAD(P)H, and in orange those producing ATP; metabolites (dots) that constitute biosynthetic precursors are highlighted in red. Some representative genes and their corresponding genetic interactions are included (color code of genetic interactions as previous figures).

In summary, we showed how distinct functional structures within the metabolic system, i.e., biosynthesis and catabolism, determine both the architecture of the network and its rewiring (Fig. 7C-D), an interpretation that is naturally coupled to the two main sources of robustness, i.e, redundancies and distributed compensation [27]. These predictions are based on global features of metabolism, what overcomes the limitations associated to FBA modeling (that sometimes generates artifacts due, for instance, to its latent optimality assumptions [8], but that nevertheless can provide useful conceptual guidelines to the associated biology [27]). Differential network mapping should thus consider the specific mechanistic causes of robustness in the system under study to accurately interpret the dynamic rewiring of genetic networks in health and disease [33, 34].

## Materials and Methods

### Genotypes and Simulations

We considered the iND750 genome-scale model as the WT genotype [35]. This model incorporates all the necessary complexity of *Saccharomyces cerevisiae*’s metabolism (e.g., it is fully compartmentalized), has been empirically corroborated, and also reduces the computational load associated to the background analysis. We studied two types of genotype derived from this model. Single deletion genotypes were obtained by deleting each gene present in the model individually. Neutral deletion trajectories were obtained by successively deleting genes that have no effect on phenotype [30] (i.e., optimal growth does not change) until reaching 100 deleted genes. All optimizations were performed using Flux Balance Analysis (FBA) [36]. WT growth conditions correspond to glucose minimal medium and aerobiosis (glucose: 18.5 mmol gr^−1^ h^−1^, unlimited O_2_). Fluxes through all reactions in the solution of a given genotype were normalized by the amount of biomass produced. This enables the comparison of different solutions with distinct growth rates.

### Generation and processing of genetic networks

Optimal growth of all single and double deletion mutants, encompassing all nonessential genes in a given genotype, was computed using FBA [36]. The mutant/WT growth ratios obtained were used to compute the epistasis (*ε*) that incorporated a multiplicative model and posterior scaling [12]; interactions with |*ε*| < 1.1 were not considered. An additional processing was applied to the networks to simplify functional redundancies that are non informative and do not contribute to the system-level analysis discussed in the manuscript. Namely, we identified all sets of genes coding for exactly the same reactions, and excluded all but one from each set for further analysis. This simplifies positive interactions associated to subunits of the same complex (e.g., mitochondrial ATP synthase has 15 essential subunits in the model, which results in 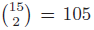 positive epistatic interactions), and also negative interactions that exist between equivalent gene duplicates coding for fitness contributing reactions. This dataset reduction was applied in all our analyses, unless otherwise specified.

### Types of genetic interactions

We classified genetic interactions in four groups according to the underlying functional structure. Synthetic lethal (SL) interactions, *ε* = −1, involve two unique alternatives for an essential function. Strong positive (SP) ones, *ε* = 1, imply an absolute functional dependence, for instance like the one found in genes that act sequentially in a parallel pathway. Two additional classes, weak positive (WP, 0.01 < *ε* < 1) and weak negative (WN, −1 < *ε* < −0.01) were defined. Weak negative interactions appear when there exists an additional less efficient solution to the two main functional alternatives represented by the WN-interacting genes. This multiplicity of alternatives with different efficiency usually reflects that (qualitatively) different ways to perform an specific function. Weak positive interactions emerge when the fitness contribution of the interacting genes is only partially overlapping, i.e., if one of the genes is deleted the other still contributes (although to a lower extent) to fitness. This could be interpreted as a form of “multifunctionality”. Note that, as positive interactions appear only among fitness contributing but nonessential genes, this implicitly reflects the presence of a less efficient and qualitatively different alternative.

### Advantages of Flux Balance Analysis

FBA was considered a suitable tool for this study due to several reasons –beyond the obvious advantage of avoiding the complications of producing the very large number of required genotypes experimentally. First, it simplifies several layers of biological complexity (e.g., gene expression or enzymatic activity regulation) by means of optimality assumptions in a relatively realistic way [36]. While this could lead to some artifacts, they do not modify in any case the general conclusions of our analysis (see also Supplement). Second, the model enables a straightforward interpretation of the phenotype as an univocal consequence of the structure of the underlying metabolic reaction network. We can thus imagine the *in silico* model as a biological “organism” *per-se*, that can provide broad conceptual guidelines for a comprehensive interpretation of the rewiring of genetic networks associated to real biological systems (not necessarily restricted to metabolism).

### Pleiotropy

We computed the pleiotropy of each nonessential and fitness contributing gene in the WT genotype following [8]. The method basically consists in optimizing for the production of a given biomass constituent individually (instead of using the entire biomass reaction) in presence and absence of a given gene. The number of affected constituents represents a rigorous measure of pleiotropy for that specific metabolic gene.

### Rewiring and metabolic modules

Background dispersion was quantified in Fig. 2B as normalized Shannon entropy *S*_MP_ (MP denotes “module pair”). This is defined as 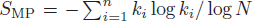, with *n* being the number of backgrounds with new interactions between the two modules, and *k_i_* the number of interactions appearing in the background *i* (divided by the total number of interactions considering all backgrounds). This was normalized by log *N*, where *N* is the total number of analyzed backgrounds where any new interaction appears between any two modules (*N* = 37). The figure illustrates how catabolic modules are characterized by appearance of many different new interactions in different backgrounds. Conversely, much fewer interactions appear among biosynthetic modules, these being generally much more background-specific. The notation of the modules in the figure can be found in Table S6.

### Random environments

1000 random environments were generated in which each of 107 organic nutrients was assigned a probability of being present from an exponential distribution (with mean = 0.1 [28]). After defining the particular set of nutrients, their dosage was randomly obtained by applying an uniform distribution between 0 and 20 mmol gr^−1^ h^−1^. All environments considered were aerobic (i.e., unconstrained O_2_ availability).

### Stability of interactions in response to environmental change

Data on instability of interactions in response to environmental change (yeast cells growing in rich media and in the presence of three distinct DNA damaging agents: Methyl methanesulfonate, Camptothecin, and Zeocin) was obtained from [17]. We considered as *not* significant epistasis those values below 2 and above −2.5 (following the original reference). We defined strong positive interactions as those in the upper quartile among positive ones, and similarly for negative ones. The instability of each category in Fig. 7A was quantified as the average number of treatments where the interaction changes or disappears (out of three). We quantified the functional similarity of the genes constituting an interaction as the ratio between the number of shared functional classes (i.e., biological process annotations as in [17]) and the minimal number of classes that one of the genes of the pair presents. If this score was bigger than 0.1 then genes were considered functionally “close”, and “distant” otherwise. In addition, we considered an interaction “stable” if it remained within the same category (sign, strength) in all conditions, and unstable otherwise (Fig. 7B).

### Robustness and genetic landscape clearly distinguish catabolic from biosynthetic modules.

We downloaded from http://www-sequence.stanford.edu/group/yeast_deletion_project/deletions3.html a list of yeast essential genes, and used the genetic interaction data of yeast metabolism from [8]. For each module, we computed the ratio between the number of essential and the number of epistatic genes, that we term *θ*. Average *θ* for catabolic and biosynthetic modules is 〈*θ*〉 = 0.18 and 0.94, respectively (Wilcoxon-Mann-Whitney test *p*=0.01), what broadly confirms the predictions of the *in silico* model (Fig. 7C).

## Acknowledgments

This research was partially supported by La Caixa Foundation PhD fellowships (CM) and the Spanish Ministerio de Economía y Competitividad BFU2011-24691 grant (JFP).

## Supplementary Tables and Figures

**Table S1.** Fold-change of metabolic production significantly correlates with rewiring, a signal that appears particularly strong in catabolic metabolites (^†^*P* < 10^−20^, ^‡^*P* < 10^−10^, ^*^*P* < 0.01). Metabolites are colored according to broad functional context where they appear. Red–catabolism; blue–biosynthesis; violet–currency metabolites and related molecules (e.g., *H*^+^ or phosphate); yellow–metabolites belonging to artifactually catabolic pathways (see below), gray–other types fo metabolites (such as water, inorganic nutrients, etc.).

**Table S2.** ATP production (*mmol grDW*^−1^ *h*^−1^) by different mechanisms in WT and in ΔCOX1 background (percentage of the total ATP produced in the cell in each case in parenthesis). FTHFL: formate tetrahydrofolate ligase, PGK: phosphoglycerate kinase, PK: pyruvate kinase. See second section Supplement for details.

**Table S3.** Distribution of genetic interactions in catabolic modules.

**Table S4.** Genetic interaction stability as a function of sign, strength and functional association.

**Table S5.** Pleiotropy is present in catabolic genes and absent in biosynthetic ones.

**Table S6.** Notation of metabolic modules associated to Fig. 2, main manuscript, that is also used trhoughout the supplement.

**Figure S1.** Differential stability of WT interactions with respect to whether their constituents genes belong to the same metabolic module (intra-module *vs.* inter-module; left), interaction strength (strong *vs.* weak; center) and type (positive *vs.* negative; right). For each interaction, instability was computed as the number of backgrounds where it disappears, changes sign, strength or both. In this figure, bars represent averages (*p*-values shown on top were obtained using the Wilcoxon test between each pair of groups).

**Figure S2.** Genetic hubs in the WT `iND750` metabolism of *S. cerevisiae*. We depict here the degrees of the top 25 most connected genes (this is the list of genetic hubs considered in the manuscript). Colors indicate the proportion of negative and positive interactions.

**Figure S3.** Schematic representation of the metabolic changes that occur upon `MIR1` deletion. Colors indicate whether a given flux has increased, decreased, shut off or on, or changed direction, as shown in the legend.

**Figure S4.** Rewiring of genetic interactions in `MIR1` background (Δ`MIR1`). Top: disrupted negative (yellow) and positive (blue) interactions. Node color represents the metabolic effect of the deletion on the reaction(s) associated to the node (see legend). Almost all the lost interactions involve either the background gene or genes that interact positively with it. Bottom-left: new negative interactions. Bottom-right: new positive interactions.

**Figure S5.** A) Schematic summary of metabolic and genetic changes after a background deletion. The cartoon is based on the Δ`MIR1` background, but aiming to provide a description of the general principles of genetic rewiring, and how it reflects metabolic changes. A) Upon deletion of a background gene (BG), a compensatory mechanism is activated (`C1`). This compensation usually originate side effects (in the case of `MIR1`, extrusion of malate from mitochondria) unless it is fully equivalent (e.g., duplicated genes). Additional mechanisms are thus activated to correct these secondary imbalances (`C1`, `C2`, `C3`, and `C4`, dashed lines). B) WT genetic network, with nodes associated to flux-carrying reactions colored green, and single fitness contribution as node size. BG interacts negatively (yellow) with compensatory mechanisms (`C1`, `C2`, `C3`, `C4`). Positive interactions (blue) occur either with the targets of its function (e.g., with respiration, R, as it is the ultimate target of phosphate import to mitochondria) or with genes that stop functioning when BG is compromised (generally displaying lower fitness effect than BG; SP for strong positive partner). C) In the mutated network, the target mechanism (respiration) interacts positively with those genes that compensate for the BG deletion (`C1`, `C2`). These compensatory mechanisms can interact among them positively or negatively, depending on the underlying functional relationship. Some of them can have in turn their own compensating mechanisms in the new background (e.g., C^′^1 buffers `C1`) also producing new negative interactions.

**Figure S6.** Rewiring of hubs MIR1 (A) and LPD1 (B) in all backgrounds. Figures equivalent to Fig. 4A, main text.

**Figure S7.** Rewiring of hubs GCV3 (A) and SER2 (B) in all backgrounds. Figures equivalent to Fig. 4A, main text.

**Figure S8.** Rewiring of hubs PMP1 (A) and ATP8 (B) in all backgrounds. Figures equivalent to Fig. 4A, main text.

**Figure S9.** Rewiring of hubs COX1 (A) and PGI1 (B) in all backgrounds. Figures equivalent to Fig. 4A, main text.

**Figure S10.** Rewiring of hubs TPI1 (A), FBA1 (B) and IDP1 (C) in all backgrounds. Figures equivalent to Fig. 4A, main text.

**Figure S11.** Rewiring of hubs ALT2 (A) and AAT1 (B) in all backgrounds. Figures equivalent to Fig. 4A, main text.

**Figure S12.** Rewiring of hubs FUM1 (A) and ACH1 (B) in all backgrounds. Figures equivalent to Fig. 4A, main text.

**Figure S13.** Rewiring of hubs RPE1 (A) and ALD6 (B) in all backgrounds. Figures equivalent to Fig. 4A, main text.

**Figure S14.** Rewiring of hubs GLY1 (A), SDH2 (B), IDP1(C) and IRC7 (D) in all backgrounds. Figures equivalent to Fig. 4A, main text.

**Figure S15.** A) Distribution of the connectivity (degree) exhibited by each hub in each background. Red dots show the WT degree, thick gray lines illustrate percentile 10 to 90, and dashed lines represent the range from the minimum (nonzero) to the maximum degree values observed. B) Number of gained and (number of) conserved genetic interactions are negatively correlated in hubs (each dot represents the Spearman’s correlation coefficient between lost WT interactions and newly gained ones; red color indicates *P* <0.001 significance). C) We classified the type of rewiring as (*horizontal*: 23 hubs, *vertical*: 37 backgrounds where they become unstable): “Collapse”, if the hub lost all its interactions; “Loss”, if it lost only part; “Partial recovery” if part of the missing genetic interactions were substituted by new ones; “Dynamic rewiring” if all of the lost interactions were substituted by new ones, so that the new degree is at least equal to that of the WT; “Gain” if the hub conserves the same partners as in the WT but gained additional ones. Moreover, “deleted” means that the node is the same as the background, and “conserved” means that the interactions of the node remain the same. Background names are colored black if they are as well hubs, red if they are not hubs in the WT but become a hub in at least one background, and gray if neither of these definitions apply.

**Figure S16.** Distribution of the number of nodes and interactions, by interaction type, in the 200 genetic networks resulting from neutral deletion accumulation trajectories. Red triangles indicate WT values.

**Figure S17.** A) Average number of negative (N) and positive (P) interactions observed to disappear and appear in genetic networks in response to neutral backgrounds. B) Average number of transitions between interaction types observed when comparing these networks with the WT one. A stronger tendency to rewiring can be observed in negative interactions overall.

**Figure S18.** Differential rewiring in LG and HG backgrounds. We show (left to right): average number of nodes whose degree is conserved (all partners equal than in the WT), number of those that conserved degree but changed their partners, number of nodes that lost interactions, and number of nodes that gained interactions.

**Figure S19.** *Top:* distribution of number of interactions, number of nodes, and interactions/node for LG (red) and HG (blue) networks (neutral backgrounds). *Bottom:* as before but restricted to the subnetworks constituted by synthetic lethal (SL, left), weak negative (WN, middle-left), weak positive (WP, middle-right), and strong positive (SP, right) interactions.

**Figure S20.** Frequency of LG and HG neutral trajectories in which a given WT interaction (dot) rewires (i.e., it disappears, changes strength or sign). A) For each interaction, a *p*-value was obtained using Fisher’s exact test (comparing the number of instabilities observed in HG and LG) and correction for multiple testing using the Benjamini-Hochberg-Yekutieli procedure; those cases with *p* < 0.001 were colored in orange. B) Fraction of weak and inter-module interactions in the “LG-enriched” and “rest” subgroups.

**Figure S21.** A) Distribution of the number of new interactions by type in LG (L) and HG (H) genotypes. For reference, we plotted the distribution of interactions in the WT network as a gray histogram line. B) For all unique interactions that appear in any of these genotypes, we plot its frequency in the HG and LG classes. We colored in red those interactions significantly enriched in LG networks (individual Fisher tests, *p*-values corrected for multiple testing using the Benjamini-Hochberg-Yekutieli procedure). C) Interactions shown in red in B are represented here as a network (edge thickness is proportional to the frequency of each interaction). D) Transition map between types of interactions in LG, and E) in HG networks.

**Figure S22.** A) For each node in the WT network, we computed the frequency it disappears from the network in LG and HG genotypes (in red those nodes that are lost more frequently in LG). B) The same was done measuring the frequency at which genes become essential in LG and HG genotypes (in red those genes that become more frequently essential in LG).

**Figure S23.** Correlation between genetic rewiring in different backgrounds and their A) degree, B) total flux through the associated reactions, C) pleiotropy, and D) single-gene fitness contribution.

**Figure S24.** We selected from the high-confidence genetic interaction dataset in [8] those interactions between genes present in the `iND750` model of *S. cerevisiae*. We separated the dataset into strong positive (upper quartile among positive in strength), strong negative (idem among negative) and weak (the rest). For functional classification, we conservatively considered a gene as “catabolic” if it belongs to either glycolysis, TCA cycle or oxidative phosphorylation; the rest were considered “biosynthetic”. According to this classification, an interaction can be catabolic, biosynthetic, or mixed (when one gene is catabolic and the other biosynthetic). In each strength category, we computed the percentage belonging to each functional class (vertical lines). To assess if any class is particularly enriched or depleted, we performed a bootstrapping analysis by randomizing 10000 times the assignment between strength and functional category of interactions. The distribution of percentages is shown in histograms, with the corresponding *p*-values. Plot boxes were colored red if significant depletion, and green if significant enrichment.

**Figure S25.** We computed, for all interactions appearing in at least one treatment, but not in the untreated network (i.e., treatment-specific interactions), the proportion of each strength/sign category. Treatment specific interactions are enriched in weak (especially weak positive) and depleted in strong (especially, in strong negative).

**Figure S26.** Genetic interactions between three genes representative of pentose phosphate pathway (ZWF1), glycolysis (PGI1) and TCA cycle (LPD1) in genotypes resulting from HG and LG trajectories. Each row represents the interactions associated to one genotype; in both cases the bottom row shows the epistasis in the WT for reference. It can be observed that the WT structure is conserved in a significant proportion of HG genotypes. In contrast, LG genotypes show a) less conservation of the WT pattern, b) generally more epistasis than in the WT, and c) a predominance of negative interactions, all of them patterns also significant at the statistical level (main text).

**Figure S27.** Synthetic lethal cluster 1.

**Figure S28.** Alternative pathways for synthesis of Ala, Ser, Gly and Trp. Gray arrows indicate fluxes in the WT metabolism. Blue arrows indicate fluxes that appear when gene names colored in blue are deleted; analogously, green arrows are fluxes that activate upon deletion of the gene colored in green.

**Figure S29.** Synthetic lethal cluster 2.

**Figure S30.** Synthetic lethal cluster 3.

**Figure S31.** Synthetic lethal cluster 4.

**Figure S32.** Synthetic lethal cluster 5.

**Figure S33.** Synthetic lethal cluster 6.

## References

1. Baryshnikova A, Costanzo M, Myers CL, Andrews B, Boone C (2013) Genetic interaction networks: toward an understanding of heritability. Annual review of genomics and human genetics 14: 111– 33.

2. Levy SF, Siegal ML (2008) Network Hubs Buffer Environmental Variation in Saccharomyces cerevisiae. PLoS Biology 6: e264.

3. Snitkin ES, Segrè D (2011) Epistatic Interaction Maps Relative to Multiple Metabolic Phenotypes. PLoS genetics 7: e1001294.

4. Roguev A, Bandyopadhyay S, Zofall M, Zhang K, Fischer T, et al. (2008) Conservation and rewiring of functional modules revealed by an epistasis map in fission yeast. Science 322: 405–10.

5. Costanzo M, Baryshnikova A, Bellay J, Kim Y, Spear ED, et al. (2010) The genetic landscape of a cell. Science 327: 425–31.

6. Reed JL, Famili I, Thiele I, Palsson BO (2006) Towards multidimensional genome annotation. Nature reviews Genetics 7: 130–41.

7. Snitkin ES, Dudley AM, Janse DM, Wong K, Church GM, et al. (2008) Model-driven analysis of experimentally determined growth phenotypes for 465 yeast gene deletion mutants under 16 different conditions. Genome biology 9: R140.

8. Szappanos B, Kovács K, Szamecz B, Honti F, Costanzo M, et al. (2011) An integrated approach to characterize genetic interaction networks in yeast metabolism. Nature genetics 43: 656–62.

9. Papp B, Pál C, Hurst LD (2004) Metabolic network analysis of the causes and evolution of enzyme dispensability in yeast. Nature 429: 661–4.

10. St Onge RP, Mani R, Oh J, Proctor M, Fung E, et al. (2007) Systematic pathway analysis using high-resolution fitness profiling of combinatorial gene deletions. Nature genetics 39: 199–206.

11. Lehner B, Crombie C, Tischler J, Fortunato A, Fraser AG (2006) Systematic mapping of genetic interactions in Caenorhabditis elegans identifies common modifiers of diverse signaling pathways. Nature genetics 38: 896–903.

12. Segrè D, Deluna A, Church GM, Kishony R (2005) Modular epistasis in yeast metabolism. Nature genetics 37: 77–83.

13. Poyatos JF (2011) The balance of weak and strong interactions in genetic networks. PloS one 6: e14598.

14. You L, Yin J (2002) Dependence of epistasis on environment and mutation severity as revealed by in silico mutagenesis of phage T7. Genetics 160: 1273–81.

15. Harrison R, Papp B, Csaba P, Oliver SG, Delneri D (2007) Plasticity of genetic interactions in metabolic networks of yeast. Proceedings of the National Academy of Sciences of the United States of America 104: 2307–2312.

16. Bandyopadhyay S, Mehta M, Kuo D, Sung Mk, Chuang R, et al. (2010) Rewiring of Genetic Networks in Response to DNA Damage. Science : 1385–1389.

17. Guénolé A, Srivas R, Vreeken K, Wang ZZ, Wang S, et al. (2013) Dissection of DNA damage responses using multiconditional genetic interaction maps. Molecular cell 49: 346–58.

18. Dixon SJ, Fedyshyn Y, Koh JLY, Prasad TSK, Chahwan C, et al. (2008) Significant conservation of synthetic lethal genetic interaction networks between distantly related eukaryotes. Proceedings of the National Academy of Sciences of the United States of America 105: 16653–8.

19. Frost A, Elgort MG, Brandman O, Ives C, Collins SR, et al. (2012) Functional repurposing revealed by comparing S. pombe and S. cerevisiae genetic interactions. Cell 149: 1339–52.

20. Ryan CJ, Roguev A, Patrick K, Xu J, Jahari H, et al. (2012) Hierarchical modularity and the evolution of genetic interactomes across species. Molecular cell 46: 691–704.

21. Chandler CH, Chari S, Dworkin I (2013) Does your gene need a background check? How genetic background impacts the analysis of mutations, genes, and evolution. Trends in genetics 29: 358–366.

22. Chari S, Dworkin I (2013) The Conditional Nature of Genetic Interactions: The Consequences of Wild-Type Backgrounds on Mutational Interactions in a Genome-Wide Modifier Screen. PLoS Genetics 9: e1003661.

23. Greenspan RJ (2009) Selection, gene interaction, and flexible gene networks. Cold Spring Harbor symposia on quantitative biology 74: 131–8.

24. Khan AI, Dinh DM, Schneider D, Lenski RE, Cooper TF (2011) Negative epistasis between beneficial mutations in an evolving bacterial population. Science 332: 1193–6.

25. Chou HH, Chiu HC, Delaney NF, Segrè D, Marx CJ (2011) Diminishing returns epistasis among beneficial mutations decelerates adaptation. Science 332: 1190–2.

26. Blank LM, Kuepfer L, Sauer U (2005) Large-scale 13C-flux analysis reveals mechanistic principles of metabolic network robustness to null mutations in yeast. Genome biology 6: R49.

27. Wagner A (2005) Robustness and Evolvability in Living Systems. Princeton University Press.

28. Wang Z, Zhang J (2009) Abundant indispensable redundancies in cellular metabolic networks. Genome biology and evolution 1: 23–33.

29. Fuhrer T, Sauer U (2009) Different biochemical mechanisms ensure network-wide balancing of reducing equivalents in microbial metabolism. Journal of bacteriology 191: 2112–21.

30. Pál C, Papp B, Lercher MJ, Csermely P, Oliver SG, et al. (2006) Chance and necessity in the evolution of minimal metabolic networks. Nature 440: 667–70.

31. Deutscher D, Meilijson I, Kupiec M, Ruppin E (2006) Multiple knockout analysis of genetic robustness in the yeast metabolic network. Nature genetics 38: 993–8.

32. Meiklejohn CD, Hartl DL (2002) A single mode of canalization. Trends in Ecology & Evolution 17: 468–473.

33. Ideker T, Krogan NJ (2012) Differential network biology. Molecular systems biology 8: 565.

34. Furlong LI (2013) Human diseases through the lens of network biology. Trends in genetics 29: 150–9.

35. Duarte NC, Herrgård MJ, Palsson BO (2004) Reconstruction and Validation of Saccharomyces cerevisiae iND750, a Fully Compartmentalized Genome-Scale Metabolic Model. Genome Research 14: 1298–1309.

36. Price ND, Reed JL, Palsson BO (2004) Genome-scale models of microbial cells: evaluating the consequences of constraints. Nature reviews Microbiology 2: 886–97.

